# Behavioral Markers of Essential Tremor Correlate with Distributed Alteration of Brain Functional Connectivity

**DOI:** 10.64898/2026.10.24.761519

**Authors:** F. Blondiaux Pirson, C. Kim, L. Colmant, L. Lebrun, A. Jankovski, G. Duchêne, L Dricot, B. Hanseeuw, F. Crevecoeur

## Abstract

Essential Tremor (ET) is a common motor disorder characterized by involuntary oscillations of the upper limbs during voluntary movement. Although increasing evidence implicates dysfunction of the cerebello–thalamo–cortical pathway, ET remains a diagnosis of exclusion, as no definitive biological marker currently exists. To better characterize ET, we recently hypothesized that impaired compensation of sensorimotor delays within the long-latency feedback pathways may generate the tremor-like oscillations observed in ET. This theory suggests that the mean spectral amplitude (MSA), derived from limb kinematics, may serve as a potential marker of ET. Here we investigated the relationship between MSA, standard clinical assessment (Fahn-Tolosa-Marin Tremor Rating Scale; FTM-TRS), and brain connectivity derived from resting state functional magnetic resonance imaging (rs-fMRI). Using seed-based analyses, we replicated previous findings of reduced functional connectivity between the dentate nucleus and components of the cerebello–thalamo–cortical pathway in ET patients compared with neurologically intact (NI) age-matched participants. Graph-theoretical analyses of whole-brain functional networks revealed distributed alterations in network topology across motor-related cortical and subcortical regions. Several graph metrics correlated with FTM–TRS scores and/or MSA, particularly within motor-related cortical and subcortical circuits, including the thalamus, basal ganglia, and cerebellum in ET. These findings indicate that ET is characterized by distributed alterations in functional brain networks beyond reduced connectivity of dentate nucleus and highlights MSA as a candidate quantitative marker linking altered brain connectivity to sensorimotor delay-driven tremor oscillations.

**Significance Statement:** How tremor severity relates to large-scale brain network organization in essential tremor (ET) remains incompletely understood. By combining quantitative tremor kinematics with resting-state functional MRI, this study links clinical manifestations of ET to alterations in both dentate nucleus functional connectivity and whole-brain network connectivity, compared with neurologically intact participants and with the conventional clinical assessment, the Fahn-Tolosa-Marin Tremor Rating Scale (FTM-TRS). Our findings indicate that spectral amplitude derived from limb oscillations in ET is associated with neural connectivity alterations, supporting its potential as a quantitative marker of disease severity. Furthermore, we identified a distributed motor-related network associated with tremor expression, providing new insights into the network-level mechanisms underlying ET.

## Introduction

Essential Tremor (ET) is a common neurological disorder affecting about 1% of the general population (Louis & Ferreira, 2010) and more than 6% of the elderly (Benito-León et al., 2003). It is typically characterized as an action tremor, with little or no tremor present at rest, while voluntary movements elicit oscillations of the limbs. It primarily affects the upper limbs and, less frequently, the head and voice (Clark & Louis, 2018; Gruol et al., 2016). Despite its high prevalence, the definitive diagnosis of ET has remained debated owing to the clinical heterogeneity of the condition (Bhatia et al., 2018; Louis, 2020b). Current clinical assessment relies largely on subjective rating scales (e.g. Fahn-Tolosa-Marin Tremor Rating Scale, FTM-TRS), and on differentiation from other well-characterized tremor disorders such as Parkinson’s disease (Algarni and Fasano, 2018). Although diagnosis remains challenging, there is growing consensus on the neurogenic origin of ET, with increasing evidence implicating the cerebello-thalamo-cortical pathway (Nicoletti et al., 2015; Tikoo et al., 2020; Holtbernd & Shah, 2021).

A recent computational framework proposes that upper-limb ET originates from a temporal mismatch in the neural mechanisms compensating for long-latency sensorimotor delays (Blondiaux Pirson et al., 2024). This explanation is based on two assumptions: (1) long-latency transcortical feedback loops incorporate a *state estimator,* i.e. a process that estimates the instantaneous state of the limb while compensating for neural transmission delays of approximately 50–60ms affecting this transcortical pathway (Crevecoeur & Scott, 2013); (2) the delay parameter implemented by the motor control system in ET may differ from the true physiological delay (Crevecoeur & Gevers, 2019). Under this framework, a mismatch between the actual and estimated delay leads to mis-timed corrective motor commands. Rather than effectively counteracting ongoing movement, these delayed corrections can partially reinforce it, producing limb oscillations. Computational simulations based on this mechanism predict exaggerated long-latency motor response to mechanical perturbations and perturbation-related oscillations whose amplitude scales with perturbation magnitude. Both features have been observed in ET.

This model predicts that the power around tremor frequency, which quantifies the amplitude of the oscillations, should increase with the mismatch between the actual and estimated delays. The framework may also provide an interpretation of the degenerative aspect of the tremor, as it captures the observation that oscillation amplitude increases over time while the dominant tremor frequency varies less comparatively. This property was also observed in the model. In this context, disease progression may reflect a gradual increase in delay mismatch, leading to a corresponding increase in tremor severity.

Here we set out to test these predictions experimentally through two objectives. First, we collected clinical and behavioral data from ET patients and neurologically intact (NI) participants. Tremor severity was assessed using the standard FTM-TRS scale and the mean spectral amplitude (MSA) derived from limb kinematics, to evaluate whether MSA could provide a complementary biomarker of tremor severity and disease stage. Second, we investigated the neural basis of ET using resting state functional magnetic resonance imaging (rs-fMRI), aiming both to replicate previous findings and to identify additional neural correlates in the mesoscale graph topology of participants’ connectomes (Benito-León et al., 2015; Fang et al., 2016; Gallea et al., 2015; Tikoo et al., 2020; Passaretti et al., 2024).

Our behavioral results revealed a positive correlation between FTM-TRS and MSA. We also replicated previous observations of altered functional connectivity between the dentate nucleus (DN) and thalamus in ET using seed-based analyses (Tikoo et al., 2020), supporting the involvement of cerebellar output in tremor generation. Graph-theoretic analyses revealed broader and more subtle alterations in whole-brain network topology in ET, extending beyond the dentate nucleus, some of which exhibited correlations with MSA or FTM-TRS. Together, these findings support the hypothesis that MSA constitutes a meaningful marker of ET and identify additional candidate regions potentially related to the tremor.

## Materials and Methods

### Participants

A total of 20 Essential Tremor (ET) patients and 22 Neurologically Intact (NI) volunteers of similar demographic repartition were enrolled in this study. All participants provided informed consent following standard procedures approved by the ethics committee of the host institution (Comité d’Éthique Hospitalo-Facultaire, UCLouvain, Belgium). ET participants who received surgical treatment for their tremor or ET participants with co-existing neurological disorders were not enrolled in the study. The control group consisted of participants who self-reported the absence of any known neurological disorders. Due to incidental findings, we excluded three participants of this group of NI volunteers. One of them presented a leukoaraiosis, and the two other excluded participants had a mega-cisterna magna. One ET participant was excluded from the study because of important artifacts in the images caused by ferromagnetic surgical material in the patient’s jaw. Analyses were therefore performed on 19 ET patients (11 females; age: M: 61,8 ± SD: 11,9) and 19 NI volunteers (11 females; age: 64 ± 14).

### Tremor Assessment

Participants were evaluated using the Fahn-Tolosa-Marin Tremor Rating Scale (FTM-TRS) (Fahn et al., 1988), which consists in a questionnaire tailored to quantify the tremor. During the FTM-TRS evaluation, participants wore wearable Inertial Measurements Units (device described in El Khoury et al., 2021), composed of accelerometers attached to the tip of their index fingers. We collected accelerometer signals attached to the fingertips in four distinct seated postures: (1) arms outstretched horizontally, with participants pointing forward with their index fingers; (2) the same arm posture but with their hands open; (3) arms approximately horizontal with the elbows flexed, such that their index fingers were pointing towards each other in front of the nose, without touching either the nose or each other, with eyes open (“finger-to-finger” posture); and (4) the same finger-to-finger posture performed with eyes closed. Each position was maintained for approximately 10 seconds, and the time windows considered were extracted manually to include only the phases during which participants had reached an approximately stable position. The accelerometers were used to acquire an acceleration signal as image of their tremor, based on which we could compute the frequency and spectral amplitude of limb oscillations based on Fourier transform of the limb kinematics (Figure 1a). Tremor frequency was estimated by treating the spectral amplitude distribution as a probability distribution and computing its expected value (Figure 1b). This definition was chosen to avoid identifying the dominant frequency from a single peak, as variability in the data often resulted in multiple peaks in the spectrum. We also extracted the MSA of the oscillations between 4 and 12 Hz (Grey area in Figure 1b). We then averaged these measurements across left and right arms and across the different postures as they did not differ significantly. This acquisition was performed on the same day as the neuroimaging assessment.

**Figure 1.**
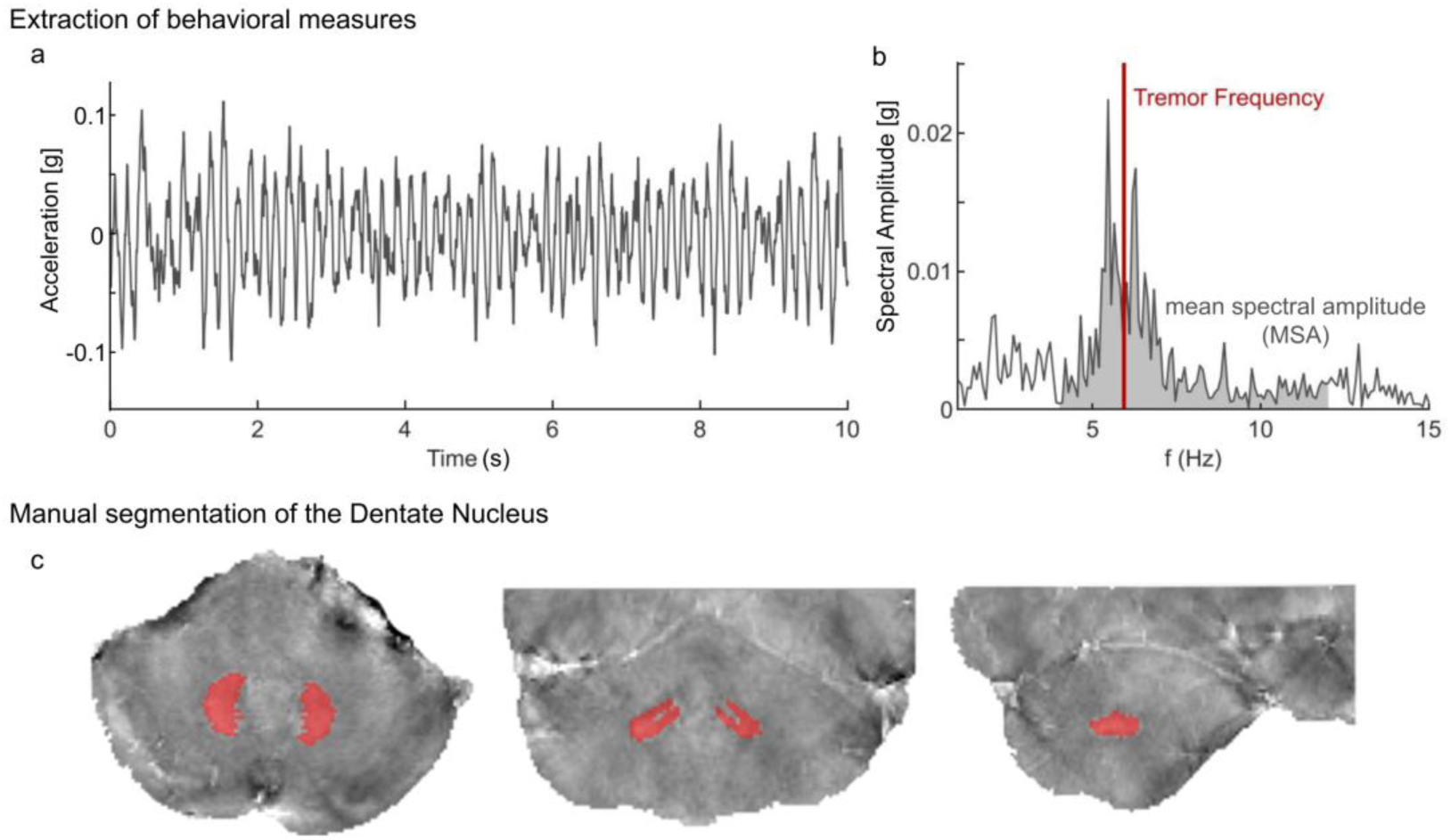
Extraction of mean spectral amplitude (MSA) and manual segmentation of the dentate nucleus. (a) Principal component of the accelerometer signals. (b) Spectral amplitude (SA) of the principal component. The grey area corresponds to the area under the curve used to estimate the MSA between 4 and 12Hz. Red vertical line corresponds to the tremor frequency computed as the expected value of the SA considered as a density function. (c) Manual segmentation of the Dentate Nucleus on the susceptibility maps.

### Neuroimaging Assessment

MRI acquisition was performed at the Cliniques Universitaires Saint-Luc (UCLouvain, Belgium) using a 3.0 Tesla head scanner (*Signa Premier^TM^* , General Electric Healthcare, United States) equipped with a 48-channel coil. ET patients and NI volunteers were positioned supine in the scanner, cushions were used to enhance comfort and to limit movement artifacts, and we used earplugs and noise-reduction headphones. Participants were instructed to remain eyes closed for the resting state as well as not to engage in cognitive or motor activities.

### A three-dimensional (3D) T1-weighted

data set encompassing the whole brain was selected to provide detailed anatomy (1 mm³) thanks to MPRAGE sequence: inversion time = 900 ms, repetition time (TR) = 2188.16 ms, echo time (TE) = 2.96 ms, flip angle (FA) = 8°, field of view (FOV) = 256*256 mm², matrix size = 256*256, 156 slices, slice thickness = 1 mm, no gap, total scan time = 5 min 36s.

### Resting state MRI T2*-weighted

sequences of brain activity was collected with echo-planar imaging: TE = 30 ms, TR = 1700 ms, FA = 90°, FOV = 220*220 mm², matrix size = 110*110, 75 slices (order ascending and interleaved), slice thickness = 2 mm, parallel imaging (Acceleration factor = 2) and hyperband factor = 3. The whole brain slices were scanned 217 times (=6 min 09s).

### A three-dimensional multi-echo gradient echo (GRE)

sequence was used to obtain T2*-weighted images. These images were used to delineate DN. Imaging parameters for the multi-echo GRE sequence were the following: TR=47.1 ms, echoes = 16, first echo time =2.76 ms, echo time spacing = 2.78 ms, bandwidth = 651.02 Hz/pixel, flip angle = 12°, FOV = 240 x 240 mm², matrix size = 256 x 256, resolution = 0.94 x 0.94 x 1.0 mm³, slices = 76, acceleration factor = 2, and total acquisition time = 5 min and 32 s. Slices in the multiecho GRE sequence were oriented axially and centered on cerebellum.

### Neuroimaging Data Processing

We analyzed fMRI data using BrainVoyager (Version 22.2.2, Brain Innovation, Maastricht, Netherlands). Preprocessing of the resting-state data consisted of linear trend removal to exclude scanner-related signal drift, a temporal high-pass filter to remove frequencies lower than 0.005 Hz and correction for head movements using a rigid body algorithm for rotating and translating each functional volume in 3D space. Additionally, we dropped the initial 7 volumes. The data was also corrected for time differences in the acquisition of the different slices and movement corrections were optimized using a sinc interpolation. As spontaneous low-frequency fluctuations can arise from non-neural signals and not only from BOLD-related fluctuations, we performed additional pre-processing steps to remove undesirable sources of variance. Regression analyses were performed to remove artifacts due to residual motions (using the six movement regressors obtained during the previous motion correction) and changes in the ventricles (the signal from the ventricular mask defined in each participant). To perform comparisons between subjects, the data was spatially normalized in the Montreal Neurological Institute (MNI) space. All co-registrations were manually verified. The data were smoothed in the spatial domain (Gaussian filter, FWHM – 5 mm). The whole brain was segmented using Freesurfer 7.0 image analysis pipeline (https://surfer.nmr.mgh.harvard.edu/). The resulting segmentation is based both on subject-independent probabilistic atlas and subject-specific measured values. We performed a visual inspection of the segmentation results and manual editing was made when needed. We only found and corrected minor skull strip errors before running the data through the pipeline again. In addition to whole brain segmentation, we also ran automatic estimation of the volume and cortical thickness of the regions defined by the Brainnetome atlas (Fan et al., 2016), which we also used for the functional connectivity analyses. We used a residual correction approach and corrected the results of the segmentation for intracranial volume (ICV), age and gender by fitting a linear regression model on all participants:

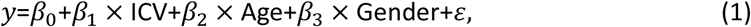

with *β*_0_ the intercept; *β*_1_, *β*_2_*, β*_3_, the coefficients associated with the independent predictors, and *ε* the residuals that captures the difference between predicted and actual values of *y*; *y* being the volume or the cortical thickness of a region of the Brainnetome or the CERES segmentation. The analyses were performed on the residuals.

Due to their higher iron content, DNs show enhanced contrast on magnetic susceptibility weighted sequences (Langkammer et al., 2012). In particular, Quantitative Susceptibility Mapping (QSM) is considered as an accurate method for DN delineation (He et al., 2017; Jäschke et al., 2023). We used the SuscEptibility mapping PIpeline tool for phAse images (SEPIA) pipeline to reconstruct QSM maps from the 3D multi-echo GRE phase data (Chan & Marques, 2021). Bilateral masks of the Dentate Nucleus were drawn manually in the QSM images using ITK-SNAP (www.itksnap.org) (Yushkevich et al., 2006) and verified by a neurosurgeon (A.J.). Segmentation for one NI participant is displayed on Figure 1c. Cerebellar lobe volumes were estimated using the CEREbellum Segmentation (CERES) pipeline. This patch-based multi-atlas segmentation tool (Romero et al., 2017) divides the cerebellum into 12 lobules in the native space. Bilateral volume of each lobule was extracted from the results.

In the seed-based analyses, the dentate nucleus was treated as a single integrated seed to capture its role as the principal cerebellar output structure and to maximize signal-to-noise ratio in this small deep nucleus. This approach also follows previous studies on this region’s connectivity to assess reproducibility of previous works (Tikoo et al., 2020). In contrast, for graph-theoretical analyses, the left and right dentate nuclei were treated as separate nodes to preserve node independence and allow potential hemispheric differences in network topology to be examined.

### Functional Connectivity Analyses

#### Seed-based analyses

Three seeds were compared for this analysis. 1) The DN as defined in the SUIT cerebellar atlas (Diedrichsen, 2006), using Left and Right dentate merged as a single region, 2) two spheres of 4-mm radius centered on DN following the SUIT atlas. Coordinates in the MNI space for the left and right DN were respectively (-18, -58, -36) and (18, -56, -36). The two masks were merged to obtain a single bilateral mask of the DN. This seed reproduces the work from Tikoo and colleagues (Tikoo et al., 2020)., 3) manual masks of the DN for each individual. We constructed for each subject three correlation maps using the three seed definition. Using these maps, group comparisons between NI and ET were investigated via voxel-wise t-tests. We studied correlations between functional connectivity maps and the FTM-TRS score in all participants via voxel-wise correlation tests. For the ET subgroup we also considered the other relevant behavioral measures (tremor frequency and spectral amplitude). P-values were corrected against multiple comparisons using a False Discovery Rate (FDR) of 0.01. Figure 3 present results using the seed as defined in the SUIT atlas.

### Graph theoretical analyses

#### Parcellation and network construction

To balance spatial resolution and statistical power, we analyzed two brain parcellation schemes of different sizes. 1) a moderate-resolution network (127 nodes) was defined using the Hammers atlas, which includes 95 regions (Hammers et al., 2003). We replaced the two cerebellar regions by a more detailed parcellation of the cerebellum using the SUIT cerebellar atlas comprising 34 cerebellar regions (Diedrichsen, 2006), and 2) a higher-resolution network (280 nodes) was constructed using the Brainnetome atlas, which parcellates the bilateral hemispheres into 246 regions (Fan et al., 2016), again combined with the same SUIT cerebellar atlas. We used data prior to spatial smoothing to generate these matrices. Matrices were generated by computing the bivariate Pearsons’s correlation between the time series of the extracted mean BOLD signal for each pair of nodes, resulting in a 127 x 127 and 280 x 280 connectivity matrix per individual. We converted the resulting coefficients to normally distributed scores using Fisher’s transformation. This transformation improves the normality assumptions of the second-level analysis. The 127-node network was used to investigate the functional connectivity between the broader brain regions, whereas the 280-node network was analyzed exploratorily to compare with the findings with those from the seed-based analysis, with the results reported in the Supplementary Materials (Tables S2, S3, and S5). Given the relatively small sample size (19 individuals with essential tremor and 19 NI volunteers), the smaller parcellation allows for less stringent FDR correction, while still preserving sufficient anatomical specificity to interpret regional network alterations (Fornito et al., 2010; Zalesky et al., 2012). Conversely, the higher resolution parcellation provides greater anatomical specificity and may reveal changes in connectivity in subregions that could be obscured when signals are averaged across larger parcels.

#### Thresholding

To ensure comparability across subjects, density-based thresholding was applied to each connectivity matrix. To reduce variability associated with selecting a single arbitrary threshold, we computed the area under the curve (AUC) of each graph metric across a range of network densities, providing a threshold-independent evaluation of network topology (Lü, L. and Zhou, T., 2011; Zhu et al.,2017). For each subject, negative correlations were first set to zero prior to density thresholding, as negative weights are commonly excluded in functional connectivity graph construction due to their ambiguous interpretation in network topology (Rubinov & Sporns 2010; Zalesky et al., 2012). The strongest connections were retained until a target network density was reached, after which the resulting adjacency matrices were binarized for graph-theoretical analyses.

To determine the minimum and maximum density (ρ) for AUC calculation, binarized matrices were examined across densities ranging from 10% to 80% (increment = 1%, i.e., 0.01). The minimum density was set to 0.23 (23%), at which all networks across all participants were sufficiently connected (>85% of nodes in the networks were connected). The upper density limit was set to 0.45 (45%) where across-subject variance of global clustering remained stable since densities above ρ ≈ 0.50 showed substantial fluctuations, suggesting a possible influence of noisy or unreliable edges. Accordingly, the AUC of graph metrics was computed as the integral of variable of interest between the density range 0.23–0.45, ensuring adequate network connectedness while preserving reliability and sensitivity to between-group differences.

#### Graph theoretical measures

Graph-theoretical measures were computed to characterize both local and global properties of the brain functional network using NetworkX (version 3.5) implemented in Python (Hagberg et al., 2008). The following node-wise metrics were derived for each region: betweenness centrality (BC), closeness centrality (CC), average shortest path length (SPL), degree (D), clustering coefficient (C), triangle (T), and local efficiency (LE). These metrics were selected because they are commonly used in graph-theoretical analyses and collectively capture complementary aspects of network organization, despite some mathematical dependencies between individual measures (e.g., SPL and CC). Metrics describing network integration (BC, CC, and SPL) capture the efficiency of information transfer within the network, whereas metrics describing segregation and local connectivity (C, T, and LE) characterize the organization and robustness of local neighborhoods. D may contribute to both local connectivity and global network integration.

Standard graph-theoretical definitions and formulas for these metrics are summarized below. BC quantifies how often a node *i* lies on the shortest paths between all other pairs of nodes in the network. Regions with high BC may act as communication hubs that facilitate information transfer between different brain regions:

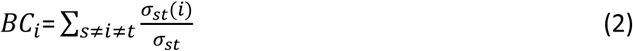

where *σ_st_* denotes the total number of shortest paths from node *s* to node *t*, and *σ_st_*(*i*) represents the number of those shortest paths that pass through node *i*. SPL of node *i* quantifies the mean number of steps required to reach all other and lower SPL values indicate more efficient information transfer and greater network integration:

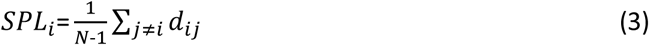

where *N* is the total number of nodes in the network, and *d_ij_* denotes the shortest path (number of edges) between *i* and *j*. CC measures how close a single node *i* is, on average, to all other nodes via shortest paths. Higher CC values indicate that a region has more efficient access to other regions through relatively short paths:

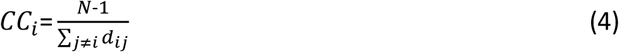

T represents the number of triangles involving node *i* and reflects the tendency of neighboring brain regions to form tightly interconnected local circuits:

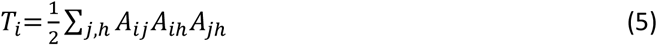

where *A_ij_* denotes the binary adjacency matrix. D is defined as:

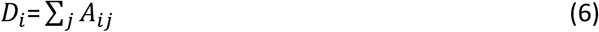

where *A_ij_* denotes the binary adjacency matrix. Nodes with higher D are more strongly connected to other brain regions and may play an important role in network integration. C quantifies the extent to which the neighbors of node *i* are interconnected. Higher C values reflect stronger local specialization and functional segregation:

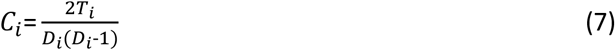

where *T_i_* denotes the number of triangles involving node *i* (Eq. 5), and *D_i_* represents the degree of node *i* (Eq. 6). LE quantifies local fault tolerance and resilience of the network surrounding node *i*. It is defined as the average inverse shortest-path length between the neighbors of node *i* after removal of node *i*. Higher LE values indicate that neighbouring nodes can maintain efficient communication through short alternative paths despite the removal of node *i*:

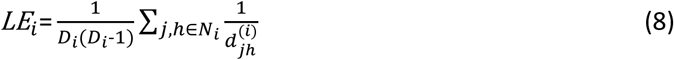

where *N_i_* denotes the set of neighbours of node *i*, *D_i_* represents the degree of node *i* (Eq. 6), and *d_jh_^(i)^_jh_* denotes the shortest path between nodes *j* and ℎ in the subgraph obtained after removing node *i*.

#### Statistical analysis on graph metrics

Node-wise AUC of graph metrics, computed across a density-based threshold range (0.23–0.45), was used for group difference and associations with behavioral measures (FTM-TRS and MSA). Group differences in graph-theoretical measures were assessed using analysis of covariance (ANCOVA), with group as the between-subject factor and age and gender included as covariates. Statistical analyses were performed in Python using the Pingouin package. To obtain robust statistical inference given the modest sample size, permutation-based testing was implemented using the Freedman–Lane procedure with 5,000 permutations within the ANCOVA framework (Freedman & Lane, 1983; Winkler et al., 2014). This approach provides *p*-values without relying on strong distributional assumptions and is well suited to analyses involving covariates and relatively small samples. To control for multiple node-wise comparisons, resulting *p*-values were adjusted using the Benjamini–Hochberg procedure to control FDR, providing a balance between Type I error control and statistical power (Benjamini & Hochberg, 1995)

Both uncorrected (p < 0.05) and FDR-corrected (q < 0.05) values are reported. Multiple comparisons were performed across 127 (280) regions, and as expected, FDR correction substantially reduced the number of statistically significant findings. We therefore evaluated both sets of results; FDR-corrected regions as robust findings and the uncorrected results as exploratory. We note however that some regions exhibited both group differences and significant correlations with behavioral measures. These two tests being independent we can estimate that in these cases, the rate of false-positive for these results was in fact much less than ∼5% and potentially closer to 0.25% (p ∼ (0.05)^2^). Thus, although exploratory, the corresponding uncorrected results can be considered reliable.

To examine the association between brain network properties and behavioral measures, partial correlation analyses were performed between node-wise AUC values of graph metrics and two behavioral measures (FTM-TRS and MSA). Age and sex were included as covariates in all analyses (Cohen et al., 2003). Covariate effects were regressed out from both graph metrics and behavioral measures, and Pearson correlation coefficients were computed between the resulting residuals. Statistical significance was assessed using a non-parametric permutation approach (Nichols & Holmes, 2002), in which behavioral scores were randomly permuted across subjects 5,000 times. For each permutation, the correlation coefficient was recomputed to generate a null distribution for each node, and two-sided empirical p-values were derived accordingly. FDR correction was applied across nodes using the Benjamini–Hochberg procedure; however, no associations remained significant after correction in 127-node network. Therefore, these findings are considered exploratory, but findings are reported using uncorrected permutation p-values (*p* < 0.05), together with the corresponding partial correlation coefficients (*r*) as complementary effect-size measures to characterize the magnitude and direction of the observed associations that may warrant further investigation.

## Results

### Demographic and behavioral measures

There were no significant differences in age (*p*=0.61, *t*_36_=0.52) or gender distribution (p=1, *t*_36_=0) between the two groups. ET participants had an FTM-TRS score of 21.18 ± 2.16 (mean ± SEM) while the NI participants had a score of 2.82 ± 0.59. Scores significantly differed between groups (*p*<10^-4^, *t*_36_=-8.2). ET participants had a tremor frequency of 6.33 ± 0.27 Hz. We calculated correlations between essential tremor frequency and behavioural measures (Figure 2a–c). Notably, significant correlations were observed between the FTM-TRS clinical scale and the MSA at the tremor frequency (Figure 2b), indicating a positive association between tremor severity and limb kinematics in ET. In contrast, the corresponding correlation was significantly weaker in the NI group than in the ET group (independent-samples *t*-test, two-sided, *p* = 0.0015).

**Figure 2.**
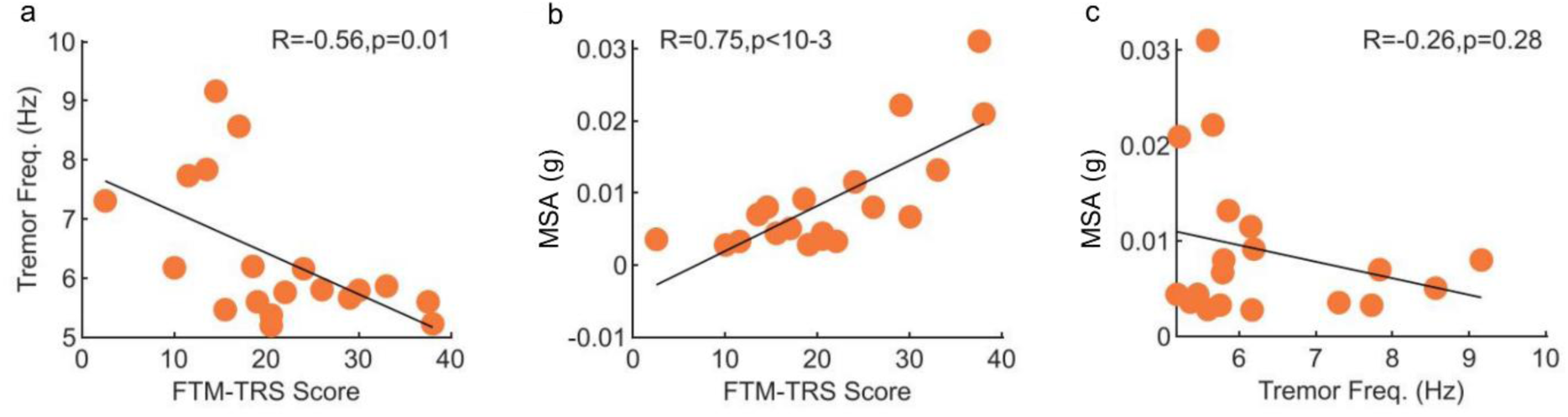
Correlation between behavior measures. (a) Correlation between Tremor Frequency and FTM-TRS Score. (b) Correlation between MSA and FTM-TRS Score. (c) Correlation between MSA and Tremor Frequency. Abbreviations: FTM–TRS, Fahn-Tolosa-Marin Tremor Rating Scale; MSA, mean spectral amplitude.

### Volumetric Analyses

#### Dentate Nucleus

Using QSM images, we manually segmented the DN of our participants, an exemplar NI participant is displayed on Figure 1c. We did not find significant differences of volume between groups (t-test: *t*_36_=0.21 , =0084)), eiiherr a crrellatinn iihh hee remorr ccree Peaason’ss correlation: 0.16, *p*=0.32).

#### Cerebellum

No significant changes in cerebellar volume were found between the two groups. A significant negative correlation was found between the FTM-TRS score and the volume of the lobule X (r=-0.4, p-uncorrected=0.014, adjusted for multiple comparisons using FDR (q = 0.2)). Results for all lobules are available in Supplementary Table S1.

#### Brainnetome Regions

No significant changes of cortical thickness were found between the two groups neither significant correlation between the cortical thickness and any of the behavioral measures.

### Seed-based analysis of the DN functional connectivity

We first aimed at reproducing previous results on functional connectivity of the dentate nucleus (DN) using a seed-based approach as reported in previous work (Tikoo et al., 2020). We observed a decreased functional connectivity in ET patients between the DN and the bilateral thalamus, bilateral basal ganglia, supplementary motor area, right primary motor area, right temporal lobe (inferior and transverse temporal gyrus), left precuneus and left superior frontal gyrus (Figure 3a; regions in red). We found increased functional connectivity in the ET group between the DN and the cerebellar lobes and the medulla oblongata (Figure 3a; regions in blue). Finally, we found a negative association of the FTM-TRS score in the thalamus (Figure 3b). The MSA and the tremor frequency were not associated with significant changes in DN functional connectivity. All cluster coordinates and sizes are reported on Table 1.

**Figure 3.**
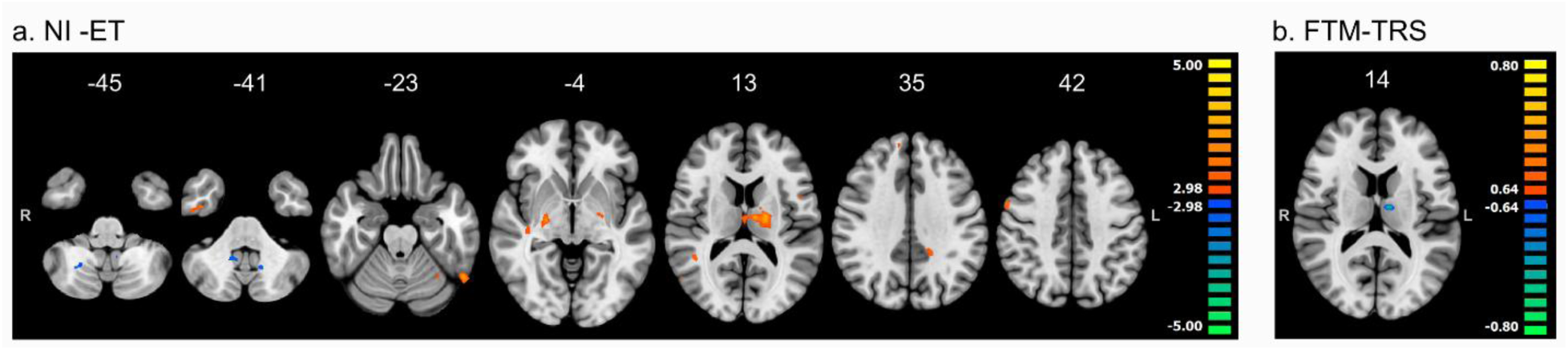
Dentate nucleus (DN) functional connectivity. (a) Group differences in the DN functional connectivity between ET and NI individuals. Maps of significant differences superimposed on T1 MNI template, red-yellow clusters imply higher DN functional connectivity for NI participants, blue-green clusters imply higher DN functional connectivity for ET patients (two-sample t-test, two-tailed, p-uncorrected < 0.005, FDR corrected at q = 0.01. Color bars represent t values. (b) Correlation of the DN functional connectivity with the FTM-TRS score in all participants. Blue-green clusters imply lower DN functional connectivity in participants with high FTM-TRS score (p-uncorrected *p*<10^-6^, FDR corrected at q = 0.1). Color bars represent r values. Abbreviations: ET, Essential Tremor; NI, neurologically intact; FTM-TRS, Fahn-Tolosa-Marin Tremor Rating Scale.

**Table 1.**
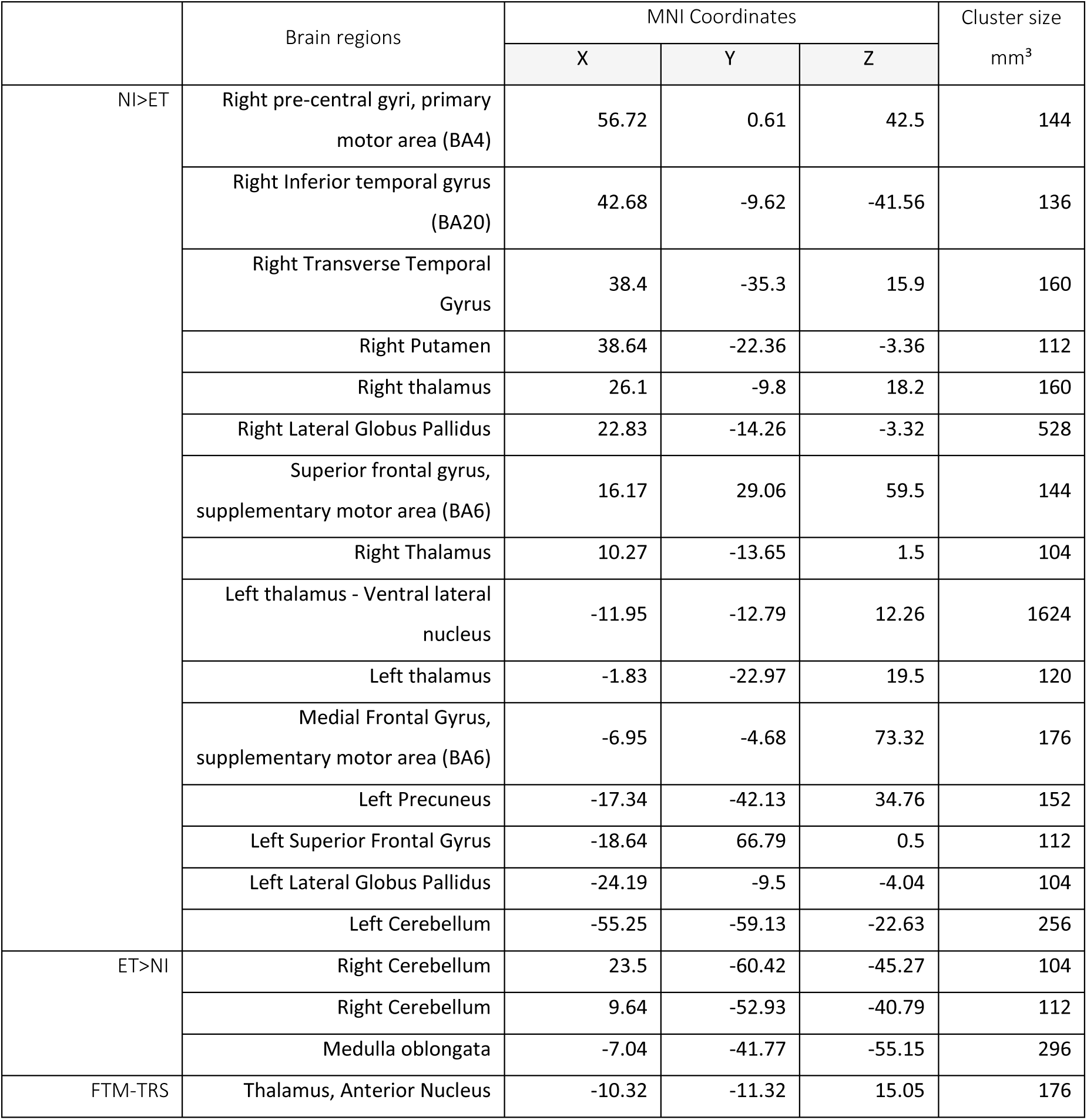
Brain regions showed significant differences of dentate nucleus (DN) functional connectivity between groups or significant correlations between DN functional connectivity and FTM-TRS. The results for group comparisons are corrected for multiple comparisons using FDR (q = 0.01) and correlations with tremor scores using FDR (q = 0.1). Cluster sizes are expressed in *mm*^3^ and coordinates are expressed in MNI152 space. Abbreviations: FTM-TRS, Fahn-Tolosa-Marin Rating Scale; BA, Brodmann Area; ET, Essential Tremor; NI, neurologically intact.

| | Brain regions | MNI Coordinates | | | Cluster size<br>$mm^3$ |
| --- | --- | --- | --- | --- | --- |
|  |  | X | Y | Z |  |
| NI>ET | Right pre-central gyri, primary motor area (BA4) | 56.72 | 0.61 | 42.5 | 144 |
|  | Right Inferior temporal gyrus (BA20) | 42.68 | -9.62 | -41.56 | 136 |
|  | Right Transverse Temporal Gyrus | 38.4 | -35.3 | 15.9 | 160 |
|  | Right Putamen | 38.64 | -22.36 | -3.36 | 112 |
|  | Right thalamus | 26.1 | -9.8 | 18.2 | 160 |
|  | Right Lateral Globus Pallidus | 22.83 | -14.26 | -3.32 | 528 |
|  | Superior frontal gyrus, supplementary motor area (BA6) | 16.17 | 29.06 | 59.5 | 144 |
|  | Right Thalamus | 10.27 | -13.65 | 1.5 | 104 |
|  | Left thalamus - Ventral lateral nucleus | -11.95 | -12.79 | 12.26 | 1624 |
|  | Left thalamus | -1.83 | -22.97 | 19.5 | 120 |
|  | Medial Frontal Gyrus, supplementary motor area (BA6) | -6.95 | -4.68 | 73.32 | 176 |
|  | Left Precuneus | -17.34 | -42.13 | 34.76 | 152 |
|  | Left Superior Frontal Gyrus | -18.64 | 66.79 | 0.5 | 112 |
|  | Left Lateral Globus Pallidus | -24.19 | -9.5 | -4.04 | 104 |
|  | Left Cerebellum | -55.25 | -59.13 | -22.63 | 256 |
| ET>NI | Right Cerebellum | 23.5 | -60.42 | -45.27 | 104 |
|  | Right Cerebellum | 9.64 | -52.93 | -40.79 | 112 |
|  | Medulla oblongata | -7.04 | -41.77 | -55.15 | 296 |
| FTM-TRS | Thalamus, Anterior Nucleus | -10.32 | -11.32 | 15.05 | 176 |

In addition, we used a more precise segmentation of DN to assess the sensitivity of the analyses across different definitions of DN. The results obtained using the DN from the SUIT atlas were compared to the results obtained by manual delineation of DN and by approximating DN as a sphere as in previous works (Tikoo et al., 2020). We did not observe any difference across these three definitions of the seed (manual drawing, sphere approximation or SUIT atlas). We computed the average t-statistic obtained when comparing NI and ET participants over all regions of the Brainnetome and SUIT atlases. We then computed the correlation between the values obtained with the three seeds and obtained a high and significant correlation for every combination (SUIT-Sphere: 0.85, *p*<10^-4^; Sphere-Manual segmentation: 0.78, *p*<10^-4^ and SUIT-Manual segmentation= 0.83, *p*<10^-4^ ) uggesstigg haat hee diffrentt aproaacess eree iiiilar. When comparing the maps of significant differences between groups obtained with the seed defined by the manual segmentation or with the seed extracted from the SUIT atlas, we observed only minor differences. To be in the best condition to study these differences, we considered a very loose uncorrected significance threshold of 0.05 and used no spatial smoothing for this analysis. These observations suggest that the different definitions of DN did not yield any statistical difference in the outcome of the seed-based analysis for our dataset.

### Graph theoretical analysis

Building on the reliable reproduction of the reduced dentate nucleus connectivity in patients with ET, as described above, we investigated differences in whole-brain connectivity constructed using two atlases of different resolutions (127- and 280-node). The brain regions showing significant differences between the ET and NI groups across graph metrics, as well as their correlations with behavioral scores (FTM-TRS and MSA), are summarized in Figure 4a and Figure 4b-c for 127-node, and Table S2-5 for 280-node, respectively. Overall, the regions identified in the group difference and the correlation with behaviour measures (FTM-TRS and MSA) were widely distributed, spanning motor-related cortical and subcortical areas. Compared to the seed-based analysis, similar brain regions were identified in the whole-brain 280-node network. Notably, the dentate nucleus was among the regions showing correlations with both behavioral measures within the 280-node network. The specific results of these analyses are described in the following sections.

**Figure 4.**
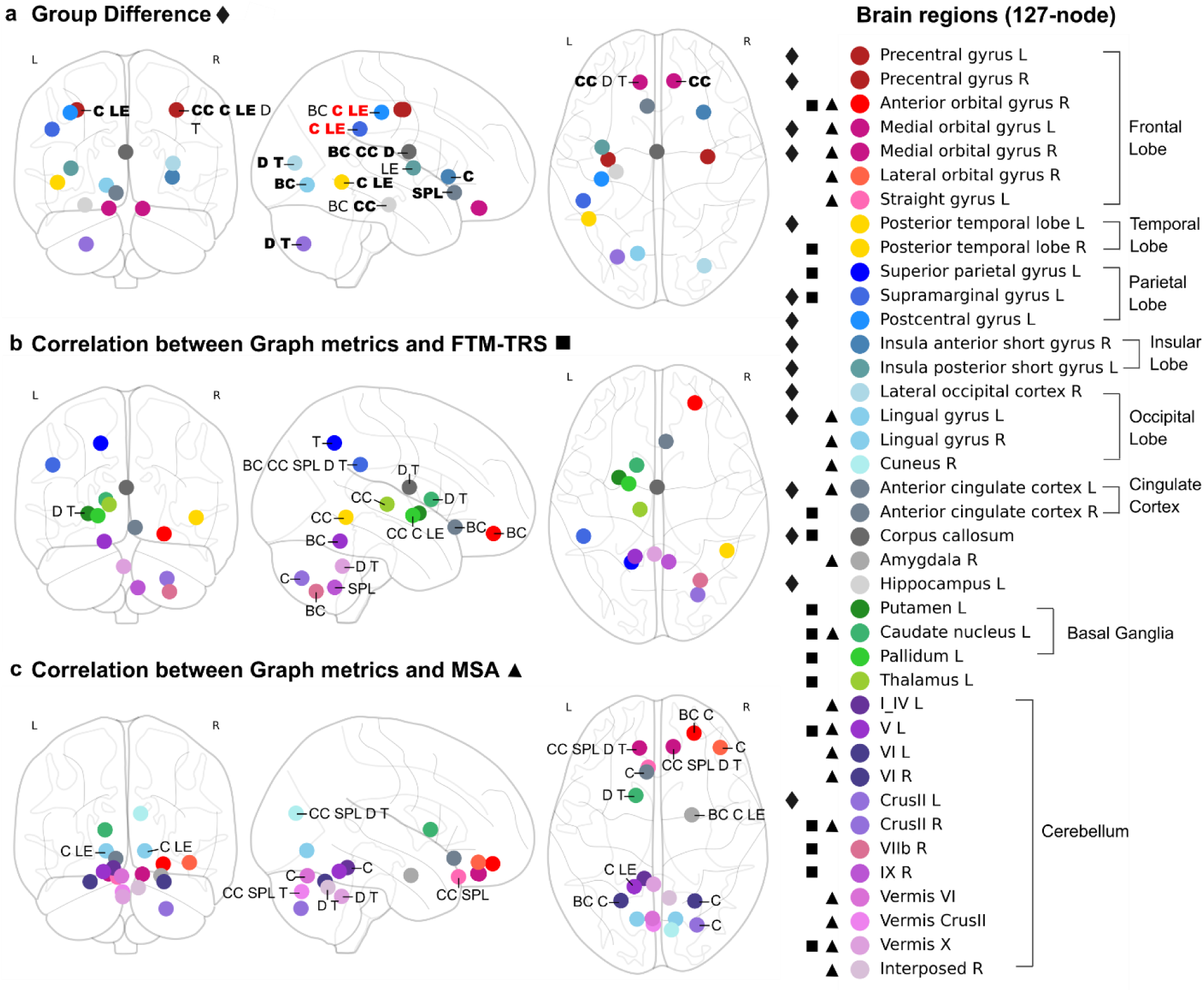
Brain regions (127-node) identified across three analyses. (a) Group difference of graph metrics between ET and NI groups. (b) Correlations between graph metrics and FTM–TRS. (c) Correlations between graph metrics and MSA. Nodes (brain regions) were identified with an uncorrected significance threshold of p < 0.05, and the corresponding brain-region names are listed in the legend according to node marker (circle) color. Graph-metric labels adjacent to node markers (circle) indicate the graph metrics in which the corresponding regions were identified. In panel (a), boldface graph-metrics labels indicate the graph metrics for which ET < NI, and graph-metrics labels in red denote those that survived FDR correction in the corresponding nodes. Nodes identified in each analysis are presented by diamonds for group difference (a), squares for correlations with FTM–TRS (b), and triangles for correlations with MSA (c). Abbreviations: FTM–TRS, Fahn-Tolosa-Marin Tremor Rating Scale; MSA, mean spectral amplitude; ET, Essential Tremor; NI, neurologically intact; BC, betweenness centrality; CC, closeness centrality; SPL, shortest path length; D, degree; T, triangle; C, clustering coefficient; LE, local efficiency.

### Group differences in graph metrics

The brain regions showing significant differences between the ET and NI groups across graph metrics are summarized in Figure 4a (127-node) and Table S2 (280-node). In Figure 4, the brain regions at an uncorrected threshold of *p* < 0.05 are shown, and graph-metric labels are highlighted in red indicate the corresponding brain regions survived FDR-correction. Significant differences were observed across multiple cortical and subcortical structures. In the frontal lobe, significant regions included the bilateral precentral gyri (primary motor cortex) and medial orbital gyri. In the temporal lobe, group differences were identified in the left posterior temporal region. In the parietal cortex, significant regions included the left supramarginal gyrus and the left postcentral gyrus (primary somatosensory cortex), while in the occipital lobe, significant effects were observed in the right lateral occipital cortex and the left lingual gyrus. Additional cortical involvement was found in the bilateral insular gyri and the left anterior cingulate cortex. Subcortical differences were identified in the corpus callosum, left hippocampus, and cerebellar regions, including the left Crus II.

Comparison with the 280-node network (Table S2), most regions identified in the primary 127-node network were also detected in the higher-resolution analysis, although they were sometimes associated with different graph metrics. The 280-node network additionally revealed more spatially specific regions within the thalamus, basal ganglia, and cerebellum, which were not detected in the 127-node network, likely due to the lower spatial resolution of the parcellation.

After applying FDR correction (q < 0.05), only a limited number of regions remained significant for the clustering coefficient and local efficiency (highlighted in red in Figure 4a): left supramarginal gyrus and the left postcentral gyrus in parietal cortex in the 127-node network, and the left superior temporal sulcus in the 280-node network (Table S3).

According to Figure 4a, where boldface metrics denote smaller values in ET relative to NI (ET < NI), the ET group exhibited predominantly reduced global integration and local connectivity across multiple cortical regions in the 127-node network. Exceptions to this pattern were observed in the left postcentral gyrus, which showed increased global metrics, and in the right precentral gyrus, left medial orbital gyrus, and left insula, where local graph metrics were elevated in ET compared with NI. Similar patterns were observed in the 280-node network (Table S2): the bilateral postcentral gyrus showed increased global metrics, whereas the left inferior frontal and parahippocampal gyri exhibited increased local metrics. In contrast, the bilateral precentral gyrus, orbital gyri, insula, nucleus accumbens, amygdala, and interposed nucleus showed increases in both global and local metrics. Region-specific results for subcortical structures, including the basal ganglia, thalamus, and cerebellum, are detailed in Table S4.

### Correlation with behavior measures

To explore the clinical and neurobiological relevance of MSA, we examined correlations between graph metrics and both MSA and the conventional clinical severity scale (FTM–TRS) in the ET group. In the 127-node network, several regions showed significant correlations with both behavioral measures (uncorrected *p* < 0.05), including the right anterior orbital gyrus, left caudate nucleus (basal ganglia), and multiple cerebellar regions (Figure 4b–c). In the 280-node network (Supplementary Table S5), a more spatially distributed pattern of associations was observed, with additional subdivisions extending from frontal cortical regions to the cerebellum, including the left dentate nucleus, which were identified in both correlation analyses. After applying FDR correction, the left inferior parietal area (A40rd) remained significant for BC in the FTM-TRS analysis, while the left precentral gyrus (A6cdl) remained significant for BC in the MSA analysis.

Regarding the MSA analysis and overlapping regions with other analysis, the brain regions correlated with MSA were primarily located within motor-related networks, including the motor-related frontal cortex, basal ganglia, and cerebellum in 127-node network. The 280-node network identified additional regions, including the thalamus and dentate nucleus. The regions shared between the group differences and the correlations with MSA in the 127-node network (Figure 4a and c) were distributed across the frontal, occipital, and cingulate cortices, as well as the cerebellum. Additional overlapping regions, including the thalamus and basal ganglia, were identified in the 280-node network (Tables S2 and S5). Overall, the regions showing overlap between the group differences and the correlation analyses with both FTM-TRS and MSA were primarily located within motor-related cortical and cerebellar circuits implicated in essential tremor. This convergence provides neuroanatomical evidence that MSA may reflect disease-related network alterations associated with tremor severity, supporting its potential relevance as a complementary biomarker.

With respect to correlation coefficient (*r*), positive correlations in global metrics (BC and CC) and negative correlation in SPL suggest increased participation of these regions in global communication pathways as severity increases. Positive correlations in local graph metrics (C, LE, and T) generally indicate alterations in local connectivity with greater tremor severity. Positive correlations in D also suggest that regions become connected to a greater number of brain regions as tremor severity increases. Overall, temporal, parietal, and occipital cortices, along with most cerebellar regions, exhibited predominantly positive correlations, whereas frontal and cingulate cortices and most subcortical regions including the basal ganglia and thalamus (except for cerebellum) showed negative correlations in both behavioral analyses (Figure 5). Especially in the MSA analysis, frontal regions demonstrated largely negative correlations across both global and local metrics, with the exception of the right anterior orbital gyrus. Within the cerebellum, most regions showed positive correlations, except for right Crus II and Vermis X, which exhibited negative associations. Notably, in the 280-node network, the left dentate nucleus exhibited positive correlations in both analyses. For the FTM-TRS associations, the correlation coefficients were *r* = 0.502 (*p* = 0.04) for C and *r* = 0.486 (*p* = 0.048) for LE, whereas for the MSA associations, they were *r* = 0.658 (*p* = 0.004) for C and *r* = 0.635 (*p* = 0.03) for LE (Table S5).

**Figure 5.**
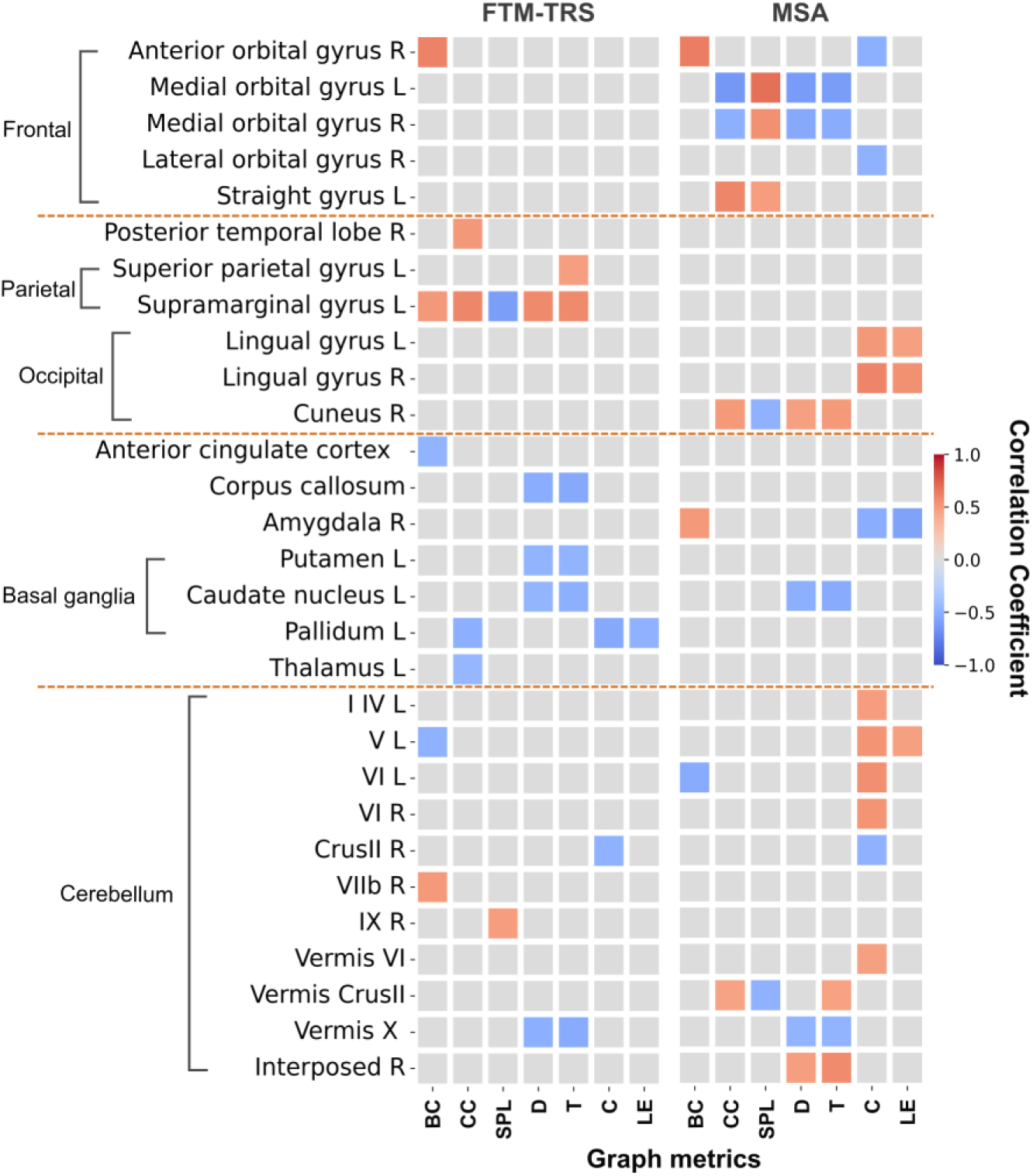
Brain regions showing significant correlations were identified using permutation tests (uncorrected p < 0.05). Red and blue colors indicate positive and negative correlation coefficients, respectively. SPL is inversely related to global network integration, with lower SPL values reflecting more long-range communication across the network. Positive correlations indicate that higher graph metric values are associated with higher behavioral scores (i.e., more severe tremor), whereas negative correlations indicate the opposite relationship. Abbreviations: FTM–TRS, Fahn-Tolosa-Marin Tremor Rating Scale; MSA, mean spectral amplitude; BC, betweenness centrality; CC, closeness centrality; SPL, shortest path length; D, degree; T, triangle; C, clustering coefficient; LE, local efficiency.

Across all analyses, an exact regional overlap was not observed; however, convergent involvement of cerebellar, thalamic, and motor-related systems was consistently identified. In line with the replication of the findings (Tikoo et al., 2020), the DN emerged as a key structure underlying brain alterations in essential tremor, with its role evident even at the level of whole-brain network topology in the 280-node network. Beyond this central role of the DN, graph-theoretical analyses further revealed more distributed alterations across motor-related cortical and subcortical networks, indicating that ET involves widespread network-level dysfunction rather than being confined to region-specific connectivity alone.

## Discussion

This study pursued two main objectives. First, we investigated whether the MSA derived from tremor kinematics from ET patients could serve as a potential biomarker for tremor severity by comparing it with the conventional clinical severity measure (FTM–TRS). Second, we examined brain network alterations in ET using rs-fMRI through both seed-based and graph theoretical analyses. In the seed-based analysis, functional connectivity of the DN was used to independently replicate previously reported alterations in dentate connectivity in ET (Tikoo et al., 2020). Beyond this region-specific analysis, we investigated whole-brain alterations in functional network topology using graph-theoretical metrics derived from functional connectivity networks constructed from 127- and 280-node brain parcellations. To further address the first objective, we examined correlations between functional connectivity measures and both behavior measures (FTM-TRS and MSA) to identify brain regions whose connectivity was associated with tremor severity, thereby providing evidence for a link between tremor kinematics at the behavioral level and their underlying neural networks.

Our seed-based analyses corroborated the findings of Tikoo and colleagues (Tikoo et al., 2020). We observed reduced DN functional connectivity with both cortical and subcortical regions, including primary motor cortex, premotor cortex, basal ganglia, and thalamus. In addition, we observed increased DN functional connectivity with cerebellar peduncles. This alteration in functional connectivity within the cerebello-thalamo-cortical pathway aligns with evidence from previous fMRI studies implicating this pathway in the pathogenesis of ET (Buijink et al.,2015; Fang et al., 2016; Holtbernd et al., 2021; Nicoletti et al., 2015). The involvement of the dentato-rubo-thamalic tract (DRTT) is further supported by the positive relationship between tremor reduction and activation of this pathway through deep brain stimulation (Middlebrooks, 2021). We also observed within our cohort of participants that a manual segmentation of the DN did not significantly influence the results of the seed-based analysis. Indeed, the maps of significant differences between groups, generated from the different seeds only marginally differed given the size and variability of our dataset.

Using a graph-theoretical framework, we further examined network-level alterations in ET compared with NI participants. The regions exhibiting altered graph metrics were largely located within motor-related cortical and subcortical circuits, including primary motor and sensorimotor cortices, basal ganglia, thalamus, and cerebellum. In general, the ET group exhibited a heterogeneous pattern of network alterations, characterized by regional differences in changes in both global and local graph metrics. Notably, local connectivity was increased within cerebellar regions, suggesting enhanced local processing within cerebellar circuits that may reflect altered sensorimotor feedback mechanisms. In parallel, increased global network metrics in cortical motor regions, such as the precentral (primary motor cortex) and postcentral gyri (somatosensory motor area), indicate enhanced long-range integration within motor networks. Together, these findings suggest that essential tremor involves a reorganization of motor-related brain networks, with strengthened local cerebellar processing alongside altered large-scale integration in cortical regions, rather than a uniform disruption of network connectivity.

When examining associations with behavioral measures, only a subset of regions overlapped between the clinical tremor scales (FTM–TRS) and kinematic measures (MSA) analyses. However, these regions were primarily located within motor-related cortical, basal ganglia, and cerebellar circuits, suggesting that both capture different but complementary aspects of tremor severity. This convergence indicates that MSA may capture aspects of tremor-related neural dysfunction, supporting its potential role as a quantitative candidate biomarker. The direction of correlation coefficients further exhibited region-specific patterns. This heterogeneous pattern may reflect differential roles of these regions within the tremor network. Negative associations observed in several subcortical structures, including the basal ganglia and thalamus, may indicate impaired coordination within motor control circuits as tremor severity increases. In contrast, positive associations observed in cerebellar regions may reflect increased recruitment of cerebellar pathways as tremor severity increases.

Interestingly, these results are in line with our previously proposed hypothesis regarding the origin of ET (Blondiaux Pirson et al., 2024). Within this framework, a computational model demonstrated that mismatches between estimated and actual sensorimotor delays give rise to tremor-like oscillations, with larger mismatches leading to increased tremor amplitude and higher MSA. Such delay compensation processes are thought to rely on the cerebello–thalamo–cortical circuit, through which error signals computed in the cerebellum are transmitted via the thalamus to primary and premotor cortices (Wolpert et al., 1998; Middleton & Strick, 2000; Shadmehr & Krakauer, 2008; Herzfeld et al., 2015). Additional support for cerebellar contribution to delay compensation comes from the studies performed with non-human primates, in which the delayed muscle responses and tremor-like oscillations emerged during cooling the cerebellum (Vilis et al., 1980; Flament et al., 1984; Hore & Vilis, 1984; Hore & Flament, 1988). Moreover, our analysis revealed associations between MSA of the tremor and functional network properties, linking tremor kinematics to underlying brain connectivity patterns. Together, these results suggest that tremor-related oscillations captured by MSA may reflect the behavioral manifestation of underlying disruptions in cerebello–thalamo–cortical network dynamics.

Our developments were based on the hypothesis that long-latency sensorimotor feedback circuits generate limb oscillations through mismatch in the delay compensation mechanism. This hypothesis justifies considering MSA as quantitative marker of the disease progression, as this variable is theoretically linked to the delay error. In addition, it also suggests that brain regions or circuits potentially altered in ET should overlap with the sensorimotor network that support long-latency feedback control. Consistent with this hypothesis, we observed substantial overlap between the main regions that exhibited differences across populations or correlations with FTM-TRS or MSA and regions involved in the long-latency sensorimotor feedback control (Scott, 2012; Kurtzer et al., 2013; Omrani et al., 2016). These regions primarily included bilateral precentral gyrus which hosts Primary Motor Cortex (group difference); parietal area 5 located in the superior parietal gyrus (correlation with FTM-TRS), as well as distributed alterations in cerebellar connectivity across several statistical comparisons (Fig. 4 and 5). Other regions located within the long-latency sensorimotor feedback networks were identified using the finer parcellation with 280 nodes, including the somatosensory cortex (Postcentral Gyrus, Table S2).

Several limitations should be considered when interpreting these findings. First, the relatively small sample size limits the statistical power of the analyses. Second, treatment withdrawal was not required prior to the assessments, which may have influenced behavioral measurements, for example by reducing tremor severity in patients receiving effective medication. Third, the cross-sectional design of the study prevents conclusions about causal relationships between network alterations and tremor severity. Finally, the large number of node-wise comparisons combined with the modest sample size resulted in only a limited number of regions remaining significant after FDR correction. Therefore, some of the exploratory findings should be interpreted cautiously and require confirmation in larger cohorts.

In conclusion, the present study revealed group differences in functional connectivity within the cerebello-thalamo-cortical circuit, as well as network alterations associated with tremor severity within this circuit. Additionally, we examined the associations between behavior measures (FTM-TRS and MSA) and brain networks. Overall, these results support the hypothesis that distributed motor-related regions including the DN play an important role in the tremor generation in ET and highlight MSA as a potential biomarker linking tremor kinematics to altered brain network organization. Future work should focus on developing more detailed models that link network topology to motor control and further clarify how alterations in brain networks give rise to motor dysfunction. Such approaches are expected to provide stronger evidence for the relationship between motor impairment and underlying brain network alterations. A further limitation of this study is that only patients with essential tremor were examined. Future studies should extend this framework to other tremor disorder to determine whether MSA can serve as a general biomarker for tremor diagnosis and characterization across different tremor syndromes.

## Supplementary

### A. Annexes

**Table S1.** Correlations between cerebellar lobes volume and FTM-TRS score, corrected for ICV, age and gender. The significance level of 0.05 was adjusted for multiple comparisons using the Benjamini-Hochberg method using FDR (q = 0.2). Significance denoted in bold. Abbreviations: FTM–TRS, Fahn-Tolosa-Marin Tremor Rating Scale; ICV, intracranial volume.

| *Region* | *R* | *uncorrected*<br>*p-value* |
| --- | --- | --- |
| *Cerebellum* | -0.23 | 0.16 |
| *I-II* | -0.11 | 0.52 |
| *III* | -0.08 | 0.62 |
| *IV* | -0.01 | 0.93 |
| *V* | 0.07 | 0.69 |
| *VI* | 0.00 | 0.98 |
| *Crus I* | -0.27 | 0.10 |
| *Crus II* | -0.02 | 0.91 |
| *VII B* | -0.20 | 0.23 |
| *VIII A* | -0.32 | 0.05 |
| *VIII B* | -0.26 | 0.12 |
| *IX* | -0.34 | 0.03 |
| *X* | **-0.40** | **0.014** |

### B. Graph theoretical analysis: Group comparison in 280-node network

The same thresholding procedure was applied to the 280-node network. The area under the curve (AUC) of graph metrics was computed across a network density range of 0.13–0.45. Group comparison results are presented in Table S2 (uncorrected p-values) and Table S3 (FDR-corrected p-values), with region names defined according to the Brainnetome and SUIT atlases. Results derived from the higher-resolution Brainnetome + SUIT atlas demonstrated convergent macro-regional patterns with those observed in the primary analyses (127-node), showing broadly similar regional distributions, with consistent involvement of motor-related cortical regions, thalamus, basal ganglia, and cerebellar regions. The detailed nomenclature of these regions is available in the Brainnetome Atlas (https://atlas.brainnetome.org/bnatlas.htm).

**Table S2.** Group comparison of AUC values (density range 0.13–0.45; uncorrected p < 0.05). Boldface metrics denote lower values in ET relative to NI (ET < NI). SPL is inversely related to global network integration, with lower values reflecting more efficient long-range communication across the network. Abbreviations: ET, Essential Tremor; NI, neurologically intact.

| Macro-Region | Left Hemisphere | Right Hemisphere |
| --- | --- | --- |
| **Betweenness Centrality** |  |  |
| Middle Frontal Gyrus | **A10l** | – |
| Inferior Temporal / Fusiform | A20rv | A20rv |
| Inferior Parietal Lobule | **A39rd** | – |
| Postcentral Gyrus (Somatosensory) | A1_2_3ulhf | – |
| Insular Gyrus | – | dld |
| Occipital Cortex | **vmPOS** | **vmPOS** |
| Cingulate Cortex | – | **A32sg** |
| Thalamus | **rTtha** | – |
| **Closeness Centrality** |  |  |
| Inferior Frontal Gyrus | A44d | – |
| Precentral Gyrus (primary motor) | A4hf | A6cdl, A4ul |
| Postcentral Gyrus (Somatosensory) | A1_2_3ulhf | A1_2_3tr |
| Orbital gyrus | A11m, A13, A12_47l | A11m |
| Superior Temporal Gyrus | A38m | – |
| Parahippocampal Gyrus | TL | – |
| Insular Gyrus | – | dld |
| Occipital Cortex | **vmPOS** | – |
| Hippocampus | rHipp | – |
| Basal Ganglia | GP, NAC | NAC |
| Amygdala | – | lAmyg |
| Cerebellum | Interposed | VIIIb |
| **Shortest Path Length** |  |  |
| Precentral Gyrus (primary motor) | – | **A6cdl, A4ul** |
| Orbital Gyrus | **A11m** | **A11m** |
| Inferior Temporal Gyrus | – | A37vl |
| Parahippocampal / Temporal Gyrus | TL | TL, A35_36r |
| Superior Parietal Lobule | A7c | A7ip |
| Inferior Parietal Lobule | A39rd | – |
| Insular Gyrus | – | **dlg** |
| Occipital / Cuneus | rCunG | rCunG |
| Occipital Cortex | mOccG, OPC, vmPOS | mOccG |
| Cingulate Cortex | A24rv | – |
| Basal Ganglia | – | vmPut |
| **Clustering Coefficient** |  |  |
| Superior Frontal Gyrus | – | **A8m, A6m** |
| Inferior Frontal Gyrus | A44v | – |
| Precentral Gyrus (primary motor) | **A4hf, A4ul** | **A6cdl, A6cvl, A4ul, A4t** |
| Paracentral Lobule | **A4ll** | **A1_2_3ll, A4ll** |
| Superior Temporal Gyrus | **A22c, A22r** | – |
| Superior Temporal Sulcus | **aSTS, rpSTS, cpSTS** | – |
| Inferior Temporal / Fusiform | **A37elv, A37lv** | **A37elv, A37lv** |
| Parahippocampal (Temporal) Gyrus | TL | – |
| Postcentral Gyrus (Somatosensory) | **A1_2_3ulhf, A2, A1_2_3tr** | **A1_2_3tr** |
| Superior Parietal Lobule | – | **A7pc** |
| Occipital Cortex | **iOccG** | **mOccG, iOccG** |
| Amygdala | – | lAmyg |
| Thalamus | **Stha** | – |
| **Local Efficiency** |  |  |
| Superior Frontal Gyrus | – | **A8m, A6m** |
| Inferior Frontal Gyrus | A44v | – |
| Precentral Gyrus (primary motor) | **A4hf, A4ul** | **A6cdl, A6cvl, A4ul, A4t** |
| Paracentral Lobule | **A4ll** | **A1_2_3ll, A4ll** |
| Superior Temporal Gyrus | **A22c** | – |
| Superior Temporal Sulcus | **aSTS, rpSTS, cpSTS** | – |
| Parahippocampal (Temporal) Gyrus | TL | – |
| Postcentral Gyrus (Somatosensory) | **A1_2_3ulhf, A2, A1_2_3tr** | **A1_2_3tr** |
| Inferior Temporal / Fusiform | **A37elv, A37lv** | **A37elv, A37lv** |
| Superior Parietal Lobule | – | **A7pc** |
| Occipital Cortex | **iOccG** | **iOccG, mOccG** |
| Amygdala | – | lAmyg |
| Basal Ganglia | vCau, NAC | – |
| Thalamus | mPFtha | – |
| Cerebellum | Vermis IX, X, Interposed | – |
| **Degree** |  |  |
| Inferior Frontal Gyrus | A44d | – |
| Precentral Gyrus (primary motor) | – | A4ul |
| Orbital Gyrus | A11m | A11m |
| Inferior Temporal Gyrus | – | **A37vl** |
| Inferior Parietal Lobule | **A39rd** | – |
| Insular Gyrus | – | dlg |
| Occipital / Cuneus | **rCunG** | **rCunG** |
| Occipital Cortex | **mOccG, msOccG, vmPOS** | **mOccG** |
| Cingulate Cortex | **A24rv** | – |
| Thalamus | **PPtha** | – |
| **Triangle Count** |  |  |
| Superior Frontal Gyrus | – | **A6dl** |
| Orbital gyrus | A11m | A11m |
| Superior Temporal Gyrus | **A41_42** | – |
| Inferior Temporal Gyrus | – | **A37vl** |
| Superior Parietal Lobule | – | **A7ip** |
| Inferior Parietal Lobule | **A39rd** | – |
| Insular Gyrus | – | dlg |
| Occipital / Cuneus | **rCunG** | **rCunG** |
| Occipital Cortex | **mOccG, msOccG, iOccG, OPC, vmPOS** | **mOccG, msOccG, OPC, vmPOS** |
| Cingulate Cortex | **A24rv** | – |
| Thalamus | **PPtha** | – |

**Table S3.** Group comparison of AUC values (density range 0.13–0.45; FDR corrected p < 0.05).

| Macro-Region | Left Hemisphere | Right Hemisphere |
| --- | --- | --- |
| **Clustering Coefficient** |  |  |
| Superior Temporal Sulcus | cpSTS | – |
| **Local Efficiency** |  |  |
| Superior Temporal Sulcus | cpSTS | – |

**Table S4.** Subcortical regions showing significant alterations in graph metrics in ET. G denotes global graph metrics, and L denotes local graph metrics. Arrows (↑, ↓) indicate the direction of change in ET relative to NI (ET vs NI).

| Basal ganglia |  | Thalamus |  | Cerebellum |  |
| --- | --- | --- | --- | --- | --- |
| nucleus accumbens (NAS) | G↑ L↑ | rostral Temporal thalamus (rTtha) | G↓ | Interposed nucleus | G↑ L↑ |
| Globus pallidus (GP) | G↑ | medial Prefrontal thalamus (mPFtha) | L↑ | Vermis IX | L↑ |
| Putamen (vmPut) | G↓ | sensory thalamus (Stha) | L↓ | X | L↑ |
| Caudate (vCau) | L↑ | posterior parietal thalamus (PPtha) | L↓ |  |  |

### C. Correlation with behavior measures in 280-node network

Correlation analyses using the high-resolution Brainnetome + SUIT atlas revealed regional patterns that were largely consistent with those observed in the primary analyses, particularly involving orbital, sensorimotor, parietal, temporal, occipital, basal ganglia, thalamic, and cerebellar regions. After applying FDR correction, the left inferior parietal area (A40rd) remained significant for betweenness centrality in the FTM-TRS analysis, while the left precentral gyrus (A6cdl) remained significant for betweenness centrality in the MSA analysis. The detailed nomenclature of these regions is available in the Brainnetome Atlas (https://atlas.brainnetome.org/bnatlas.htm).

**Table S5.**
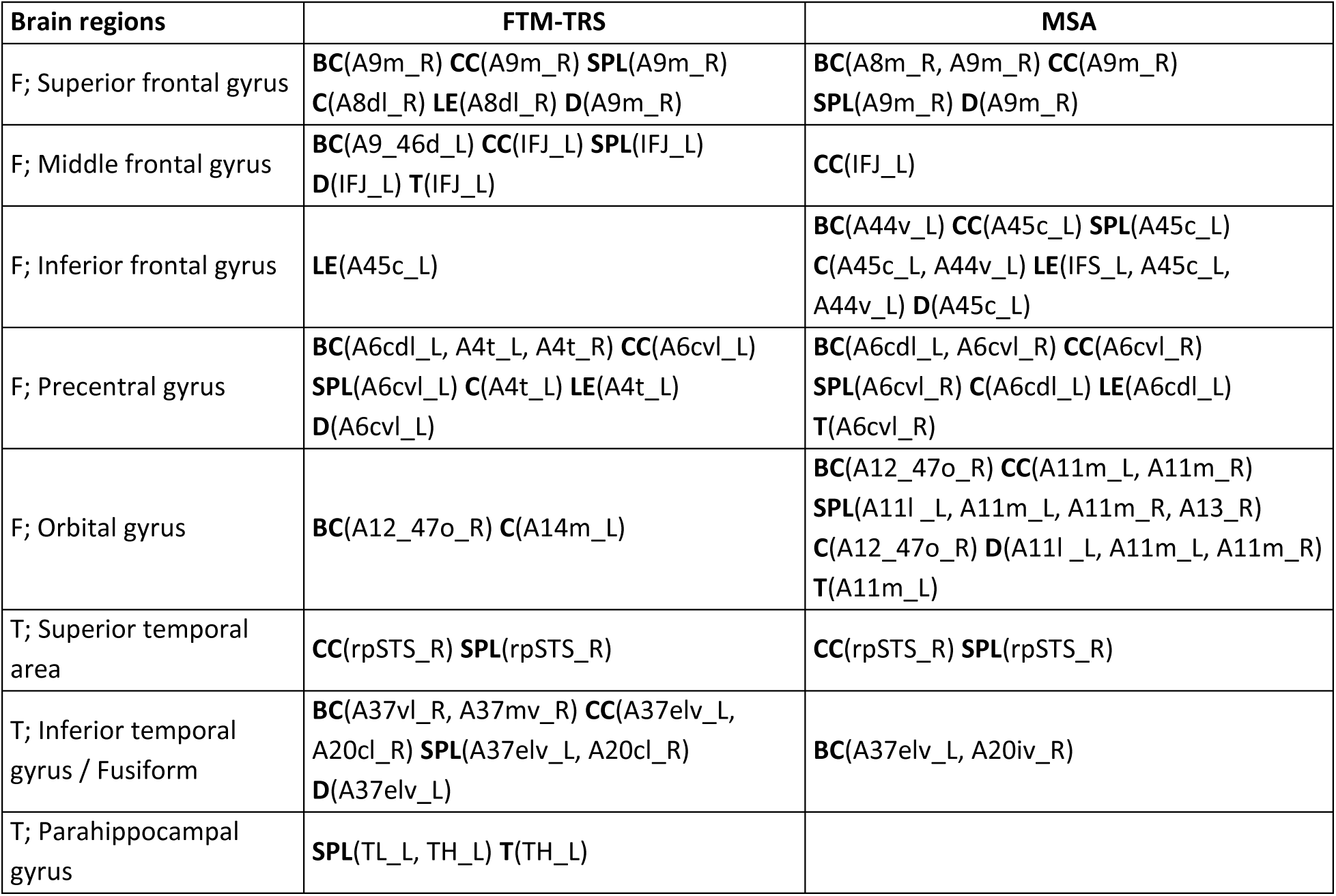
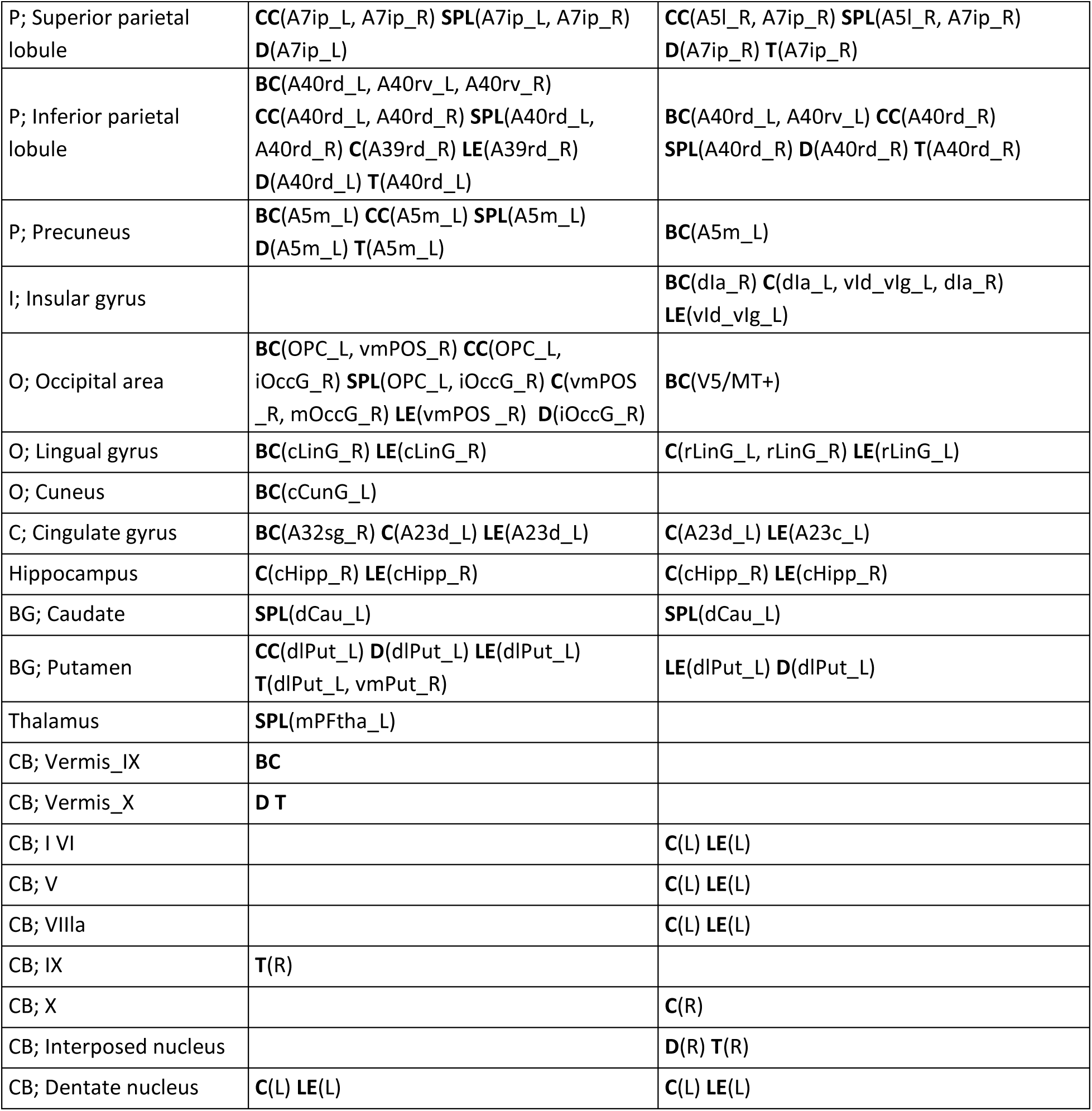
Partial correlation and permutation test between graph metrics and behavior measures (uncorrected p-value < 0.05). Abbreviations: L, left hemisphere; R, right hemisphere; F, frontal; T, temporal; P, parietal; I, insular; O, occipital; BG, basal ganglia; CB, cerebellum; FTM–TRS, Fahn-Tolosa-Marin Tremor Rating Scale; MSA, mean spectral amplitude; BC, betweenness centrality; CC, closeness centrality; SPL, shortest path length; D, degree; T, triangle; C, clustering coefficient; LE, local efficiency.

